# Thermodynamically Constrained Encoding Improves Reliability in DNA Data Storage

**DOI:** 10.64898/2026.11.13.766389

**Authors:** Jérémy Mateos, Rémi Moreau, Melpomeni Dimopoulou, Marc Antonini

## Abstract

Synthetic DNA has emerged as a promising storage medium thanks to its exceptional information density and long-term stability. However, sequence-dependent biochemical properties introduced during the encoding process can significantly affect downstream molecular processing and data recovery. Among these properties, the thermodynamic stability of DNA sequences, commonly quantified by the Folding Gibbs free energy (ΔG), can influence amplification efficiency, sequencing coverage, and ultimately decoding performance.

In this work, we experimentally investigate the impact of ΔG constraints applied during the encoding stage on DNA data recovery. We compare the reference JPEG DNA, the future image coding standard on DNA, with a ΔG-constrained variant using the same image encoded into synthetic DNA oligonucleotides. Both configurations were synthesized, amplified, sequenced, and decoded under identical experimental conditions. We analyze the relationship between predicted thermodynamic stability, sequencing coverage, and decoding performance to quantify the influence of sequence thermodynamics on the recovery of encoded information. Our results demonstrate that incorporating Δ*G* constraints during encoding substantially reduces the proportion of sequences with highly negative predicted Δ*G* values and improves sequencing coverage homogeneity, decreases the number of underrepresented oligonucleotides and dropouts, and consistently reduces the sequencing coverage required for successful decoding across three different synthesis providers, thereby reducing sequencing time. These findings provide experimental evidence that thermodynamic sequence optimization represents an effective design strategy for improving the robustness and efficiency of DNA data storage systems.

## Introduction

The exponential growth of digital data has created an increasing demand for storage technologies that combine high density, long-term durability^1,2^, and low energy consumption requirements that conventional storage media can no longer meet. Synthetic DNA has emerged as a promising alternative storage medium because of its exceptional density^3–5^, long-term chemical stability, and negligible maintenance requirements at room temperature under suitable conditions^2,6^. Theoretical estimates suggest that one gram of DNA can store exabytes of digital information making DNA an attractive candidate for long-term archival storage.

The DNA storage pipeline can be divided in 3 main stages: writing, storage, and reading. The writing process combines the encoding step where the digital data is converted into sequences composed of the four DNA nucleotides A (Adenine), C (Cytosine), G (Guanine) and T (Thymine). The encoded sequences are then synthesized into short DNA molecules known as oligonucleotides. Following storage, the oligonucleotides pass through a series of biochemical processes during the reading stage, including PCR amplification^7,8^, sequencing library preparation, and DNA sequencing. The resulting sequencing reads are then digitally processed to retrieve the original encoded digital information.

Although this general workflow has been successfully demonstrated^9–13^, several challenges remain before large-scale deployment becomes practical, with scalability representing one of the primary obstacles^14^. In addition to scalability, the reliability of the storage process is strongly affected by biochemical processes, including DNA synthesis, PCR amplification, and sequencing platforms such as Illumina^a^ and Oxford Nanopore Technology^b^. These processes introduce substitution, insertion, and deletion errors, as well as sequence-level dropouts caused by uneven amplification or sequencing bias. Consequently, DNA storage codecs aim to combine constrained sequence design with error-correcting codes to maximize decoding reliability while maintaining high storage density.

The most common biochemical constraints include controlling GC content (typically between 40% and 60%), limiting homopolymer lengths, avoiding forbidden sequence motifs, and reducing primer cross-hybridization^15,16^. These constraints are now standard components of most DNA encoding pipelines. In contrast, the use of thermodynamic constraints based on Folding Gibbs free energy (Δ*G*) during the encoding stage has received comparatively little attention and has not been extensively evaluated experimentally.

Sequences exhibiting highly negative Δ*G* values are more likely to form hairpins and self-dimers, potentially interfering with PCR amplification, and sequencing efficiency^17,18^. Although thermodynamic analysis is routinely employed for primer design and probe optimization in molecular biology, its impact when applied as an explicit encoding constraint in DNA data storage has received comparatively little experimental attention. Understanding this relationship is particularly important because uneven sequencing coverage is one of the principal causes of decoding failure. Underrepresented oligonucleotides increase the sequencing depth required for successful reconstruction and reduce the robustness of error-correction algorithms. If thermodynamic constraints can improve coverage homogeneity without significantly reducing encoding efficiency, they could provide a simple and effective strategy for increasing the reliability of DNA storage systems.

This study was motivated by observations made during the wet-lab validation of the JPEG DNA coding system. In these experiments, a header oligonucleotide was strongly underrepresented in the sequencing data, limiting successful reconstruction even though the main-section oligonucleotides presented a sufficient coverage. Because the header carries essential decoding parameters and is not protected by the Raptor redundancy (see section ‘Background’) applied to the main section, its insufficient recovery can prevent decoding of the entire file. Thermodynamic analysis of this sequence revealed a strongly negative Δ*G*, suggesting that stable secondary structures might contribute to its under representation. This observation prompted us to investigate whether thermodynamic sequence selection could improve oligonucleotide representation and decoding reliability across the encoded pool.

In this work, we experimentally evaluate the impact of introducing a thermodynamic constraint based on (Δ*G*) during the encoding stage of a DNA data storage workflow. Using the future JPEG-DNA standard^19^, we generated two encodings of the same image from the JPEG AIC-3 dataset^20^: a reference encoding and a Δ*G*-constrained encoding.

The reference configuration reused the JPEG DNA encoding previously employed for the wet-lab validation of the standard. For the Δ*G*-constrained configuration, the same source bitstream and coding parameters were retained, while additional candidate oligonucleotides were generated using the Raptor encoder. After formatting and thermodynamic evaluation, candidates with predicted folding Δ*G* values closer to zero were preferentially selected to form a decodable set containing the same number of oligonucleotides as the reference encoding.

Both configurations were synthesized independently by each of three commercial DNA synthesis providers (Agilent^c^, Dynegene^d^, and GenScript^e^) and subsequently subjected to identical amplification, sequencing, and decoding workflows, allowing the effect of thermodynamic sequence optimization to be isolated. We then investigated the relationship between predicted Δ*G*, sequencing coverage, and decoding performance across the different synthesis conditions.

Our results demonstrate that incorporating a Δ*G* constraint during encoding reduces the occurrence of thermodynamically stable secondary-structure-prone sequences, improves sequencing coverage homogeneity, decreases the number of underrepresented oligonucleotides and dropouts, and reduces the sequencing effort required for successful decoding. These improvements are consistently observed across all three synthesis providers, offering experimental evidence that thermodynamic sequence optimization constitutes an effective design strategy for improving the robustness of DNA data storage systems.

The following sections first introduce the JPEG DNA coding framework and the integration of the proposed thermodynamic constraint, then present its experimental evaluation in terms of thermodynamic properties, sequencing coverage, and decoding performance, before discussing its implications for DNA data storage.

## Background

### The JPEG DNA codec

JPEG DNA is an emerging ISO/IEC standard for storing digital media in synthetic DNA^21^. Its core coding system, ISO/IEC 25508-1, is currently at the Draft International Standard (DIS) stage^22^. The JPEG DNA encoding system consists of two main stages: source coding and channel coding. In the source-coding stage, an input image can be compressed using the JPEG XL standard^23^, producing a binary bitstream that is subsequently processed by the DNA Raptor channel encoder. However, the JPEG DNA core coding system is codec-agnostic and can therefore accommodate any source-coding method or accept arbitrary binary data directly as input to the channel encoder, independently of the JPEG XL compression stage (see Figure 1).

**Figure 1.**
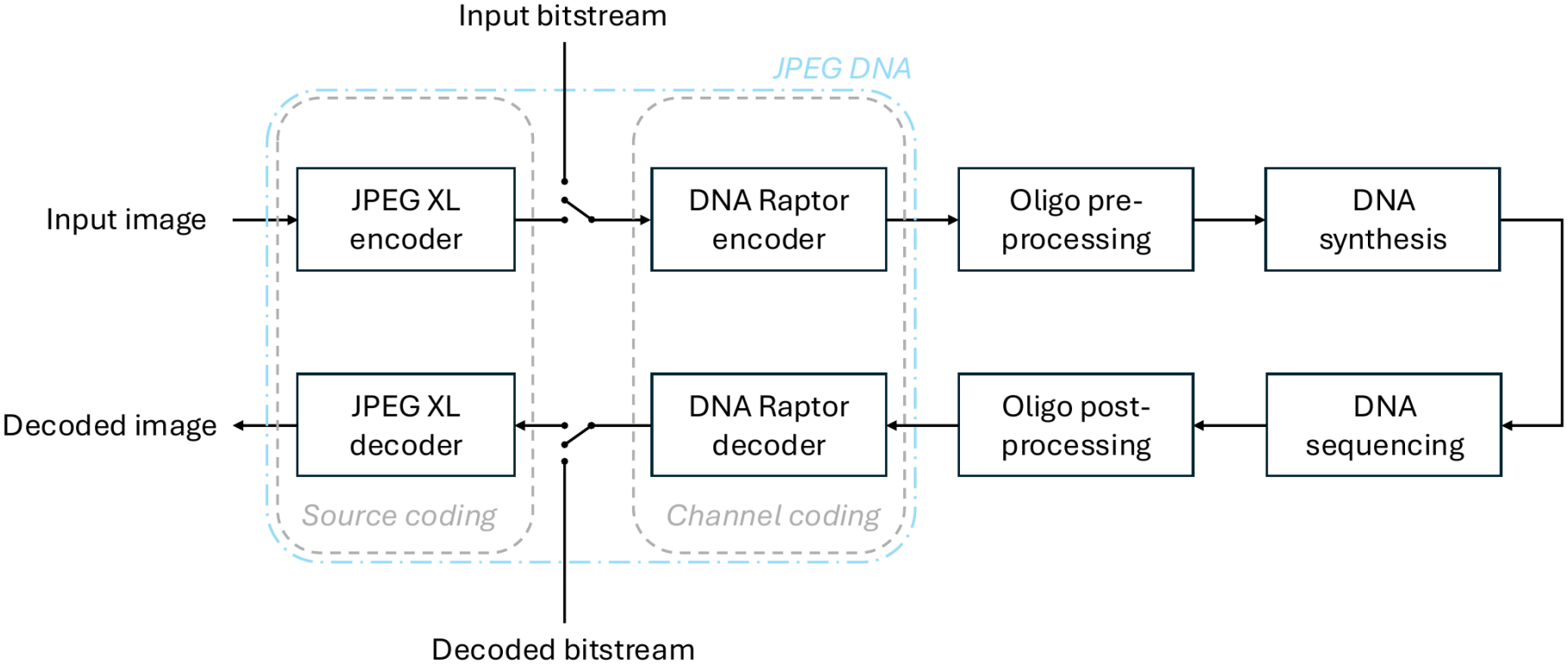
End to end workflow for digital files encoding into DNA molecules with JPEG DNA. The encoder allows images or any digital files as an input. If not an image, the JPEG XL encoder is shunted and the input bitstream goes directly inside the DNA Raptor encoder.

At the channel-coding stage, the bitstream is partitioned into source blocks and symbols then processed using a systematic Raptor code based on RFC5053^24^ specification. The resulting encoding symbols are converted into A, C, G and T sequences through a binary-to-nucleotide mapping. Since not every candidate sequence is suitable for the biochemical pipeline, each generated oligo is screened against a set of biochemical constraints. Oligos that are not respecting these constraints are discarded, and the encoding process iteratively generates new candidates until a decodable set of valid oligos is obtained. This approach, inspired by DNA Fountain codecs, takes advantage of the ability of Raptor codes to generate multiple encoding symbols from the same source data.

To provide robustness against oligonucleotide dropout, the JPEG DNA channel coding scheme introduces a coding overhead *θ* by generating additional Raptor symbols beyond the number of source symbols required to represent the data. Consequently, successful decoding does not require the recovery of every encoded oligonucleotide, provided that a sufficient number of encoding symbols is available to reconstruct the original source data, depending on the value of *θ*. Additional error protection can also be applied at the oligonucleotide level using Reed–Solomon (RS) codes to improve robustness against residual errors within individual sequences. The parameters required for decoding are stored in a separate header, which is not protected by the Raptor redundancy applied to the main section.

In the current JPEG DNA coding framework, the biochemical constraints primarily target sequence-level properties such as homopolymer length, GC content, and repeated nucleotide patterns. However, these constraints do not explicitly account for the thermodynamic propensity of an oligo to form secondary structures. In this work, we investigate the introduction of a free-energy (Δ*G*) constraint into this screening process.

## Results

### Building a thermodynamically favorable distribution using a Δ*G* constrained encoding

First, to confirm that the proposed encoding indeed generates oligonucleotides of predicted Δ*G* closer to zero, we compared two encodings of a single image from the JPEG AIC-3 dataset^20^ (`00005_560x888.png`). The reference encoding, labelled “JPEG DNA-Ref” in the figures, was compared with the Δ*G*-constrained encoding. Each configuration contained 1,983 oligonucleotides, representing the same source image with identical coding parameters but different sequence sets.

As shown in Fig. 2, applying the Δ*G* constraint substantially shifted the distribution toward less negative Δ*G* values. Because both distributions were normalized and derived from the same set of 1,983 encoded oligonucleotides, the observed shift corresponds to a redistribution of Δ*G* values rather than a change in the number of sequences. The mean predicted Δ*G* increased from −7.20 kcal/mol for the reference encoding to −4.45 kcal/mol for the Δ*G*-constrained encoding. Consequently, the proportion of oligonucleotides exhibiting highly negative Δ*G* values, which are more likely to form stable secondary structures, was markedly reduced.

**Figure 2.**
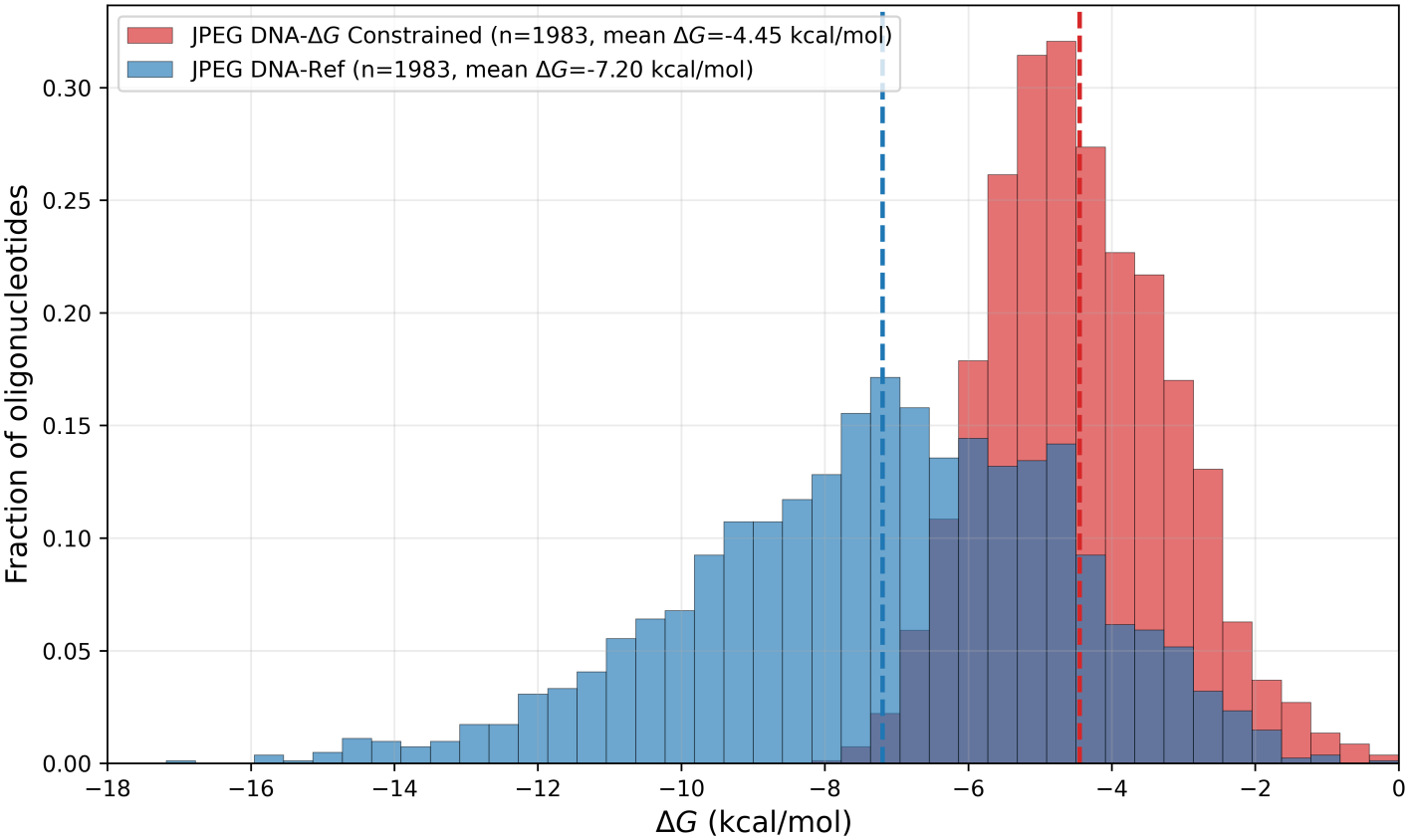
Distribution of predicted (Δ*G*) values for the reference JPEG DNA encoding and the Δ*G*-constrained encoding. Both encodings generated 1,983 oligonucleotides. Histograms are normalized to unit area, and dashed lines indicate the mean predicted Δ*G* of each distribution.

These results confirm that the proposed thermodynamic filtering effectively modifies the physicochemical properties of the encoded oligonucleotides while preserving the total number of encoded sequences. Having ensured this shift in thermodynamic stability, we next investigated its impact on sequencing coverage and decoding performance.

### Thermodynamic stability influences sequencing coverage

We next investigated whether the thermodynamic optimization introduced during encoding translated into improved sequencing representation. The sequencing dataset comprised 80,307,458 reads distributed across 13,014 FASTQ files: 25,147,839 reads for Agilent SurePrint, 31,752,670 for Dynegene, and 23,406,949 for GenScript, with 4338 FASTQ files per provider (each FASTQ file storing an average of 6171 reads). For each synthesis provider, reference coverage was normalized by the mean coverage across all 3,966 reference oligonucleotides from the two encoding configurations combined. The normalization included all references, including those with zero coverage. The mean coverage values used as normalization denominators were 4,359.40, 4,283.87, and 3,345.99 reads per reference for Agilent SurePrint, Dynegene, and GenScript, respectively. A normalized coverage of 1 therefore corresponds to the mean coverage of the combined reference and Δ*G*-constrained pools for a given provider. We first compared the normalized coverage distributions of the two encoding configurations (see Figure 3a)

**Figure 3.**
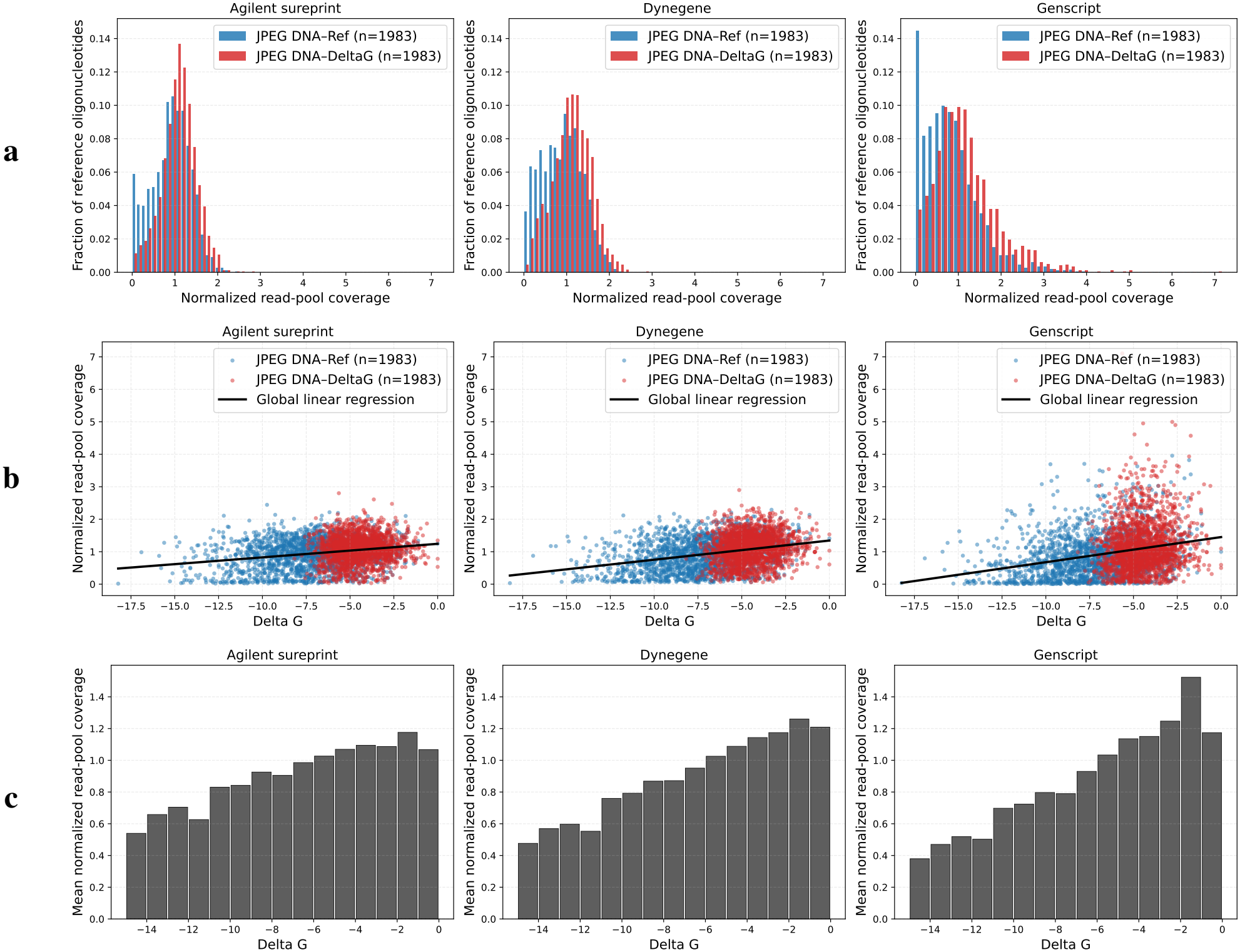
Predicted Δ*G* is positively associated with sequencing coverage across three DNA synthesis providers. **(a)**, Distribution of normalized sequencing coverage for the reference JPEG DNA encoding and the Δ*G*-constrained encoding. For each synthesis provider, reference coverage was normalized by the mean coverage across all 3,966 reference oligonucleotides from the two encoding configurations combined. The normalization included all references, including those with zero coverage. **(b)**, Normalized sequencing coverage of individual oligonucleotides as a function of predicted (Δ*G*). Black lines indicate the global linear regression fitted to both encodings, revealing a positive relationship between Δ*G* and sequencing coverage. **(c)**, Mean normalized sequencing coverage computed over successive Δ*G* intervals. Across Agilent SurePrint, Dynegene, and GenScript, oligonucleotides with less negative Δ*G* values consistently exhibit higher sequencing coverage, explaining the improved coverage homogeneity observed after Δ*G*-constrained encoding.

Across all three synthesis providers, the Δ*G*-constrained encoding produced a more concentrated distribution centered around a normalized coverage of 1, indicating a more homogeneous representation of the different oligonucleotide references.

For the reference JPEG DNA encoding, Agilent SurePrint and Dynegene exhibited relatively similar coverage distributions, whereas GenScript showed a substantially larger proportion of references with normalized coverage close to zero. Importantly, this population of strongly underrepresented references was markedly reduced by the Δ*G*-constrained encoding, including for GenScript, for which the difference was particularly pronounced.

To investigate whether these differences were associated with oligonucleotide thermodynamic properties, we analyzed the relationship between the predicted Δ*G* of each oligonucleotide and its normalized sequencing coverage (see Figure 3b). A significant positive correlation was observed for all three synthesis providers, with Spearman’s rank correlation coefficients of *ρ* = 0.195 for Agilent SurePrint, *ρ* = 0.288 for Dynegene, and *ρ* = 0.277 for GenScript (*p*-value *<* 10^*−*34^ in all cases). Oligonucleotides exhibiting less negative Δ*G* values tended to achieve higher sequencing coverage, whereas sequences with more negative Δ*G* values were more frequently associated with lower coverage.

To further characterize this relationship, oligonucleotides were grouped according to their predicted Δ*G* values, and the mean normalized sequencing coverage was calculated for each interval (see Figure 3c). The resulting profiles confirmed an overall increase in sequencing coverage as Δ*G* became less negative. This relationship was consistently observed across Agilent SurePrint, Dynegene, and GenScript, supporting a reproducible association between oligonucleotide secondary-structure stability and sequencing representation.

Together, these results indicate that reducing oligonucleotide secondary-structure stability improves sequencing representation and limits the occurrence of strongly underrepresented references. We next investigated whether these improvements translate into more efficient reference recovery and faster reconstruction of the stored payload.

### Δ*G*-constrained encoding improves decoding efficiency

Having established the association between predicted folding Δ*G* and sequencing coverage, we next investigated whether improved oligonucleotide representation translated into more efficient image reconstruction during progressive decoding.

We define progressive decoding as the process of attempting to reconstruct a file after each successive batch of sequencing reads, using all reads accumulated within the current run. For each synthesis provider, we applied this procedure to both the reference JPEG DNA encoding and the Δ*G*-constrained encoding. A run was complete once the image had been successfully reconstructed under both configurations. The next run began with the next unused FASTQ batch, so each run used a distinct set of reads. We continued until all available batches had been processed. In the figures, *n* denotes the number of completed runs for each provider.

As detailed in the Methods section (“Read processing and decoding”), sequencing reads were processed incrementally and combined into consensus sequences using quality-weighted nucleotide voting before JPEG DNA decoding. Both encoding configurations used the same consensus procedure and error-correction parameters: a Raptor overhead of 10%, 6 bytes of Reed–Solomon redundancy per main-section oligonucleotide, and 17 bytes for the header (Methods, “Stored file and codec configurations”). Consensus reconstruction combines repeated observations of each sequence, while Reed–Solomon coding provides protection against residual errors within its correction capability. Raptor redundancy enables reconstruction from a sufficient set of recovered encoding symbols, without requiring every main-section oligonucleotide to be recovered. The separate header must also be successfully decoded. A reduction in the sequencing coverage required for image reconstruction therefore does not necessarily imply that individual reads contain fewer errors or that fewer reads are needed to obtain each correct consensus. A more uniform distribution of oligonucleotide abundances can reduce the total number of sequencing reads required for successful decoding. By limiting the oversampling of highly abundant oligonucleotides and improving the representation of less abundant ones, a sufficient number of distinct oligonucleotides can be recovered at a lower sequencing depth. Therefore, this comparison specifically assesses the impact of thermodynamic sequence selection, while keeping the consensus reconstruction and error-correction procedures unchanged. The individual contributions of these two decoding steps were not evaluated separately.

Across all three synthesis providers, the Δ*G*-constrained encoding consistently reduced the sequencing coverage required for successful image reconstruction (see Figure 4a). For Agilent SurePrint, the mean coverage at successful decoding decreased from 6.063 for the reference JPEG DNA encoding to 4.431 for the Δ*G*-constrained encoding, corresponding to a 26.9% reduction. Similarly, for Dynegene, the mean coverage decreased from 6.266 to 4.732, corresponding to a 24.5% reduction. The largest improvement was observed for GenScript, for which the mean coverage at decoding decreased from 20.05 to 7.734, representing a 61.4% reduction. The higher coverage required by the GenScript reference encoding meant that more reads were consumed per progressive decoding run, resulting in fewer complete runs from the available sequencing dataset.

**Figure 4.**
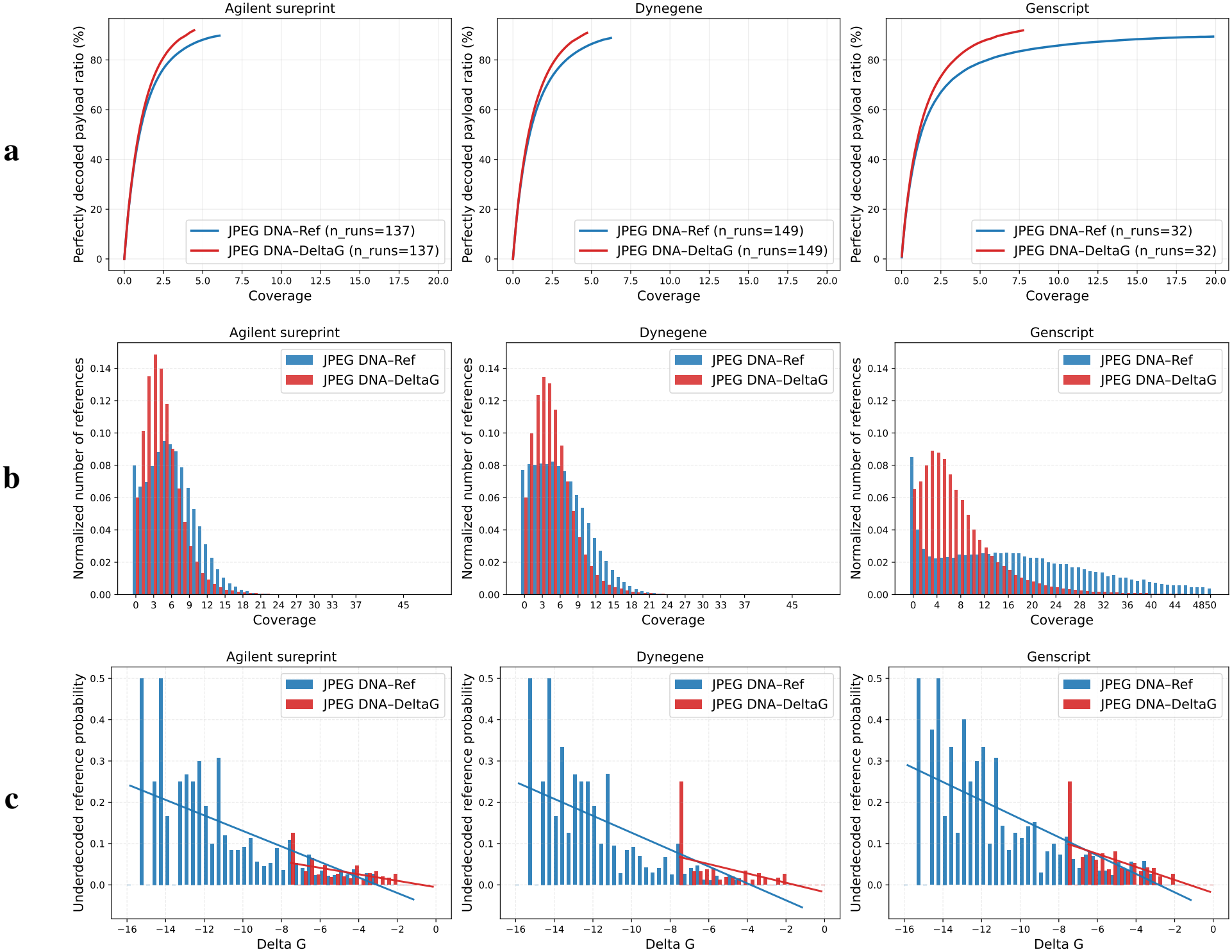
Δ*G*-constrained encoding reduces the sequencing coverage required for successful image decoding across three DNA synthesis providers. **(a)**, For each encoding configuration, the curve shows the mean percentage of reference oligonucleotides perfectly decoded, averaged across independent decoding runs. The number of runs (*n*) is given in the legend of each plot. **(b)**, Distribution of reference oligonucleotide coverage at the time of complete image decoding for the reference JPEG DNA and Δ*G*-constrained encodings. **(c)**, Probability that a reference remains underdecoded as a function of its predicted Δ*G*. A reference is considered underdecoded when no corresponding sequencing reads were available to the decoder at the end of the decoding run for most (*>* 50%) of the decoding runs.

We next examined the sequencing coverage distribution of individual reference oligonucleotides at the time of complete image decoding (see Figure 4b). Across all three synthesis providers, the Δ*G*-constrained encoding shifted the distribution of individual reference coverage toward lower values. The median reference coverage decreased from 6 to 4 for both Agilent SurePrint and Dynegene, and from 18 to 6 for GenScript. This shift was particularly pronounced for GenScript and is consistent with the lower global sequencing coverage required for successful decoding observed in Figure 4a.

To further investigate the relationship between thermodynamic stability and reference recovery, we quantified the probability that a reference remained underdecoded as a function of its predicted Δ*G* (see Figure 4c). Here, we distinguish underdecoded references from underrepresented references. An underrepresented reference is present in the sequencing data but exhibits lower coverage relative to the other references, whereas an underdecoded reference has not yet been observed by the decoder at a given stage of progressive sequencing, indicating that no corresponding sequencing read has been recovered.

The probability of remaining underdecoded increased toward more negative Δ*G* values across all three synthesis providers. A strong negative correlation was observed between Δ*G* and underdecoded probability for Agilent SurePrint (*ρ* = −0.72, *p*-value = 1.05 × 10^*−*11^), Dynegene (*ρ* = −0.76, *p*-value = 5.47 × 10^*−*14^), and GenScript (*ρ* = −0.77, *p*-value = 1.80 10^*−*14^). Therefore, oligonucleotides with more negative Δ*G* values, and consequently a greater tendency to form stable secondary structures, were more likely to remain unavailable to the decoder during progressive sequencing.

Together, these results demonstrate that thermodynamic sequence optimization is associated with improved sequencing representation and decoding performance. By reducing the occurrence of oligonucleotides with highly negative Δ*G* values, the Δ*G*-constrained encoding limits the number of underrepresented and underdecoded references, enabling successful image reconstruction at lower sequencing coverage across all three synthesis providers.

## Discussion

The results of this study demonstrate the direct impact of thermodynamic constraints on DNA encoding and subsequent data recovery. By reducing the occurrence of sequences with highly negative Δ*G* values, the constrained encoding improves sequencing coverage homogeneity and enables more efficient and faster successful decoding. To trace the source of these observed improvements, we hypothesize that thermodynamic constraints benefit biochemical processes, including DNA synthesis and PCR amplification. By limiting the formation of stable secondary structures, thermodynamic optimization may reduce sequence-dependent biases and the loss or underrepresentation of specific oligonucleotides during these processes.

### Faster decoding through improved sequencing coverage

Improving sequencing coverage directly benefits the decodability of DNA-stored data. A more homogeneous representation of the oligonucleotide pool allows the decoder to recover the information required for successful reconstruction with fewer sequencing reads. In the context of progressive sequencing, this means that the decoding threshold can be reached earlier, thereby reducing the time required to recover the stored file. The improvement provided by the Δ*G* constraint can therefore translate into faster data retrieval, which becomes particularly relevant as the volume of stored data increases. At a given sequencing depth, this more uniform coverage also reduces the number of oligonucleotides that remain unobserved, thereby limiting effective dropout and improving the robustness of decoding and data reconstruction.

### Theoretical decoding capacity of a PromethION flow cell

The experimentally determined coverage required for successful file reconstruction can be used to estimate the theoretical amount of digital data recoverable from a single PromethION flow cell. Importantly, successful JPEG DNA decoding does not require the recovery of 100% of the reference oligonucleotides, as the redundancy introduced by the Raptor code and Reed–Solomon error correction allows the original file to be reconstructed despite missing or incorrectly decoded sequences. Therefore, the relevant metric for this estimation is the sequencing coverage at which the complete file becomes successfully decodable rather than the coverage required to recover the entire oligonucleotide pool.

Assuming that we synthesize the oligo pool with Genscript such that a decoding run requires (as previously seen) an average coverage of *c* = 6.063 for the original JPEG DNA encoding (resp. 4.431 for JPEG DNA-Δ*G* constrained encoding), knowing that the original image of size *s* = 852.2 kilobytes was encoded in *n*_1_ = 1, 983 oligos, and that our promethion flowcell produced a total of *n*_tot_ = 122.477 million sequencing reads, a lower bound lb_flowcell_ on the theoretical decoding capacity (expressed in bytes) of the promethion flowcell can be estimated by the formula:

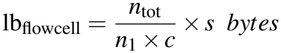

Numerically, we estimate using this formula that one flow cell can decode at least lb_flowcell_ = 8.861 gigabytes of data encoded with the original JPEG DNA algorithm, and lb_flowcell_ = 11.88 gigabytes of data encoded with its Δ*G*-constrained counterpart. The Δ*G*-constrained encoding provides a capacity increase of 37%, verifying once again that it represents an effective design strategy for improving DNA data storage efficiency.

### Codec Integration of the Δ*G* constraint

The experimental results presented in this work provide a strong motivation for integrating the Δ*G* constraint directly into the JPEG DNA encoding process. In the present study, the thermodynamic constraint was applied through a post-encoding filtering strategy: an excess of candidate oligonucleotides was first generated by the JPEG DNA codec, after which their predicted Δ*G* values were computed and sequences with more favorable thermodynamic properties were preferentially selected. While this approach enabled the experimental evaluation of the effect of thermodynamic filtering independently of the existing codec implementation, a more direct strategy would consist in incorporating the Δ*G* criterion into the biochemical constraint screening stage of the encoder itself.

The JPEG DNA encoder already iteratively generates candidate sequences and evaluates them against biochemical constraints before retaining them as valid encoding symbols. The Δ*G* constraint could therefore be introduced as an additional criterion within this existing screening process, alongside constraints on GC content, homopolymer length, and repeated nucleotide patterns. Candidate oligonucleotides exhibiting a predicted Δ*G* below a predefined threshold could be rejected, prompting the Raptor encoder to generate an alternative encoding symbol until a thermodynamically valid sequence is obtained.

This integration would eliminate the need for the post-encoding generation and filtering strategy used in the present study and would systematically enforce thermodynamic constraints during sequence generation. The results shown in this work therefore provide an experimental basis for considering thermodynamic stability as an intrinsic biochemical constraint in future implementations of the JPEG DNA codec.

### Impact of thermodynamic constraints on long-oligonucleotide synthesis

The effect of the Δ*G* constraint was particularly pronounced for the GenScript pool, as shown in Figure 4. In the present study, oligonucleotides synthesized by GenScript reached a total length of 202 nt, exceeding the standard length of 170 nt specified for its high-throughput semiconductor-based synthesis platform and requiring a specific agreement with the provider.

This observation raises the possibility that thermodynamic sequence properties become increasingly important when approaching or exceeding the nominal length range of an oligonucleotide synthesis technology. As oligonucleotide length increases, the number of possible intramolecular interactions also increases, potentially favoring the formation of stable secondary structures in sequences exhibiting highly negative Δ*G* values. Such structures may contribute to sequence-dependent biases during or following synthesis, resulting in reduced representation or loss of specific oligonucleotides from the synthesized pool. The particularly pronounced benefit of the Δ*G* constraint observed for the 202 nt GenScript pool is consistent with this hypothesis.

Although synthesis-related effects cannot be formally isolated from downstream experimental steps, the same PCR amplification, library preparation, and sequencing protocols were applied to all three synthesis providers, with samples processed in parallel within the same experimental workflow to minimize manipulation-related biases. The particularly pronounced effect observed for the GenScript pool may therefore reflect differences arising from the synthesized oligonucleotide pools themselves, although dedicated characterization of oligonucleotide abundance directly after synthesis would be required to establish this relationship.

Nevertheless, these results suggest that thermodynamic optimization may facilitate the reliable use of longer oligonucleotides, particularly when operating beyond the standard synthesis length of a given platform. As high-throughput synthesis technologies evolve toward longer DNA molecules, incorporating thermodynamic constraints at the encoding stage could therefore become increasingly important for exploiting increased strand lengths while limiting sequence-dependent biases that may compromise data recovery.

## Methods

### Filtering design

The experimental dataset was derived from the same image and reference oligonucleotide set previously used for the wet-lab validation of the JPEG DNA standard. Two encoding configurations were generated from this dataset: the original JPEG DNA encoding (refered as “JPEG DNA-Ref” in the figures) and a Δ*G*-constrained encoding. Both configurations represented the same image using 1,983 oligonucleotides and identical source-coding and error-protection parameters. They were processed using the same experimental and computational procedures to evaluate the effect of thermodynamic sequence constraint on sequencing representation and decoding performance.

### In vitro experimental workflow

#### Stored file and codec configurations

The stored file corresponds to the image `00005_560x888.png` from the JPEG AIC-3 dataset, with a file size of 856 kB. The image was encoded using the JPEG DNA codec with JPEG XL (v0.11.1) as the image compression component. A JPEG XL quality parameter of *Q* = 78 was used, corresponding to a subjective visual quality score of −0.75 Just Noticeable Difference (JND) with the original image. The JPEG DNA encoding was configured with a Raptor symbol size of 29 and a Raptor overhead *θ* of 0.1, corresponding to 10% additional symbols. For the main section, 6 bytes of Reed–Solomon redundancy were used. Since each Reed–Solomon byte is represented by 4 nt, this corresponds to 24 nt of redundancy. The resulting configuration therefore consists of a 132-nt payload followed by 24 nt of Reed–Solomon redundancy, resulting in a 156-nt encoded main section.

The JPEG DNA header is shorter than the main section. To obtain a uniform encoded sequence length, additional Reed– Solomon redundancy was used to pad the header. Specifically, the header was protected using 17 bytes of Reed–Solomon redundancy, resulting in an encoded length of 155 nt. A single `A` nucleotide was appended to match the 156-nt length of the encoded main-section oligonucleotides before the addition of tags and primer-binding regions.

#### Oligonucleotide design and synthesis

For each DNA supplier (Agilent, Dynegene, and GenScript), the encoded payload sequences were designed identically, with a final payload length of 156 nt for the main section and 155 nt for the header section, followed by a single `A` padding nucleotide.

A dataset-specific 6-nt tag to adress both the reference and the Δ*G*-constrained encoding configurations, was then appended to the 5^*′*^ end of each sequence. It is followed by the addition of a 20-nt forward primer (AGAGCGGCGTATTGTATTCG) and a 20-nt reverse primer (AGTTCACGTCCAGTCAGAGT). The primer sequences were generated using the DSP Tools software^25^.

The resulting oligonucleotides had a total length of 202 nt, exceeding the standard maximum synthesis length offered by GenScript (170 nt), for which a specific agreement was obtained. Each supplier-specific oligonucleotide pool contained both the reference and the Δ*G*-constrained encoding configurations.

#### Constraint-based filtering

For the filtered JPEG DNA DeltaG variants, an excess pool of candidate sequences was first generated using the codec JPEG DNA. The encoded payloads were then formatted to match the 202-nt oligonucleotide structure described above, including the configuration-specific tags and primers. The sequences were subsequently screened and retained only if they satisfied the following (Δ*G*, in kcal/mol) constraint computed with the NUPACK Python API^26,27^ (v4.0.1.8) with temperature parameters set at 55^*°*^C, corresponding to the primer annealing temperature (*T*_*a*_). Sequences with Δ*G* values closest to zero were preferentially selected.

#### PCR, library preparation, and sequencing

The oligo pool from Agilent, Dynegene and GenScript, received dehydrated, was resuspended to a concentration of 1 ng *µ*L^*−*1^ using nuclease-free water. For PCR amplification, 10 *µ*L of the resuspended oligo pool,corresponding to 10 ng of DNA, was mixed with 25 *µ*L KAPA HiFi HotStart ReadyMix 2 × (Roche), 1 *µ*L of 5 *µ*M forward primer, 1 *µ*L of 5 *µ*M reverse primer, and 13 *µ*L nuclease-free water, yielding a final reaction volume of 50 *µ*L. The reaction was prepared in separate tubes. Thermocycling followed established protocols, consisting of an initial denaturation at 95^*°*^C for 5 min, followed by 15 cycles of denaturation at 95^*°*^C for 20 s, annealing at 55^*°*^C for 30 s, and extension at 72^*°*^C for 20 s. PCR products were subsequently purified using SPRI magnetic beads (Beckman Coulter) and analyzed on a BioAnalyzer system using the High Sensitivity DNA kit and reagents (Agilent Technologies).

Libraries for Oxford Nanopore sequencing were prepared using the SQK-NBD114.24 Native barcoding kit (Oxford Nanopore Technologies). Nanopore sequencing was performed on a PromethION platform using R10.4.1 PromethION flow cell.

#### Read processing and decoding

Sequencing reads are received and processed incrementally, one new batch (as a `.fastq` file) being received every minute during the sequencing. We process the reads batch per batch and decoding run per decoding run. One *decoding run* is defined by the repetition of one or more decoding passes, one per read batch, until the same image had been successfully reconstructed from both encoding configurations by the JPEG DNA algorithm. One *decoding pass* is defined by the following steps:

1. **Preprocessing**. For each read in the batch processed during this pass: locate and trim its primer and tag sequences to extract the payload in between, then allocate the read to the corresponding image. To efficiently compute the decoding runs over our 500 GB of reads, the read preprocessing and clustering steps are computed using the algorithm minimap2^28^: non-aligned reads are discarded while each aligned read is trimmed according to the computed alignment, then moved to the cluster associated to the found reference;
2. **Clustering and consensus**. For each encoding configuration, newly received payloads were assigned to their corresponding reference oligonucleotides and added to the clusters accumulated during the current run. A consensus sequence was then computed for each group using the quality-weighted voting procedure described below;
3. **Image decoding**. The consensus sequences of each configuration not yet decoded were supplied to the JPEG DNA decoder. Once the image was successfully reconstructed from a configuration, that configuration was marked as decoded and its decoding endpoint was recorded;
4. **Stopping criterion**. Once both encoding configurations had been successfully decoded, the accumulated reads and decoding state were cleared, and a new run started with the next batch. Otherwise, the current run continued for the configuration or configurations not yet decoded.

For consensus reconstruction, only payload reads matching the expected sequence length were retained. At each nucleotide position *l*, the observation from read *r* was assigned a weight derived from its Phred quality score^29^ *Q*_*r,l*_:

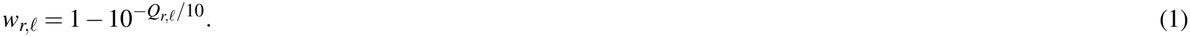

This weight represents the estimated probability of a correct base call according to the basecaller’s quality score. For a group of reads *C*, the consensus nucleotide was selected as

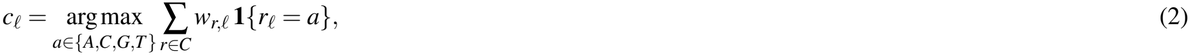

where **1** {*r*_*l*_ = *a*} equals one when read *r* contains nucleotide *a* at position *l*, and zero otherwise. The same preprocessing, consensus reconstruction, and decoding procedures were applied to both encoding configurations.

The consensus sequences were subsequently processed by the RS and Raptor decoders using the parameters specified above. RS protection corrects residual errors within its correction capability, while Raptor coding allows reconstruction from a sufficient set of recovered encoding symbols. Successful file recovery therefore does not require perfect reconstruction of every reference oligonucleotide, provided that the necessary header information is available.

Inside one decoding run, the decoding process of one image stops exactly at the first decoding pass which achieved to successfully decode it. It allows to estimate the decoding metrics of Figure 4 with precision in coverage up to the size of one batch.

Instead of the used reference-guided read clustering, a reference-blind workflow could instead combine primer and adapter trimming using Cutadapt^30^ with sequence clustering using CD-HIT^31^, as explored in related work^17^ without changing the results, since alignment is the baseline for clustering algorithm, though would have required highest computation time.

Reads and intermediate processing results were stored in a SQL database. Additional metrics were computed retrospectively, and plotting scripts retrieved the stored results to generate the figures presented in this study.

### Computational environment and software

The sequencing, along with the real-time progressive decoding runs, were computed using the Dorado basecaller on a machine equipped with four NVIDIA Ampere-series GPUs, one Turing-series GPU and 512 GB RAM.

After sequencing, all encoding and decoding computations were conducted on a personal computer equipped with an Intel Core i7-14650HX CPU (8 Performance-Cores 2.2 GHz / 5.2 GHz Turbo + 8 Efficient-Cores 1.6 GHz / 3.7 GHz Turbo - 24 Threads - Cache 30 MB) and 16 GB RAM.

## Data availability

A subset of the FASTQ files for each dataset, sufficient to decode the images synthesized by each provider, is available at Zenodo^32^. The complete FASTQ dataset and SQL databases, representing more than 500 GB of sequencing data and decoding artifacts, is available from the corresponding authors upon request.

## Acknowledgements

This work was supported totally or in part by the France 2030 investment plan, managed by the National Research Agency (ANR), through the PEPR MolecularXiv project (ANR-22-PEXM-003). The authors thank the Institute of Molecular and Cellular Pharmacology (IPMC), Sophia Antipolis, France, and its director Pascal Barbry for providing access to laboratory facilities and technical support that made the experimental work possible.

## Author contributions

J.M. designed the experiments, performed simulations, data analysis and performed the wetlab experiments. R.M. developed the implementation code and prepared the figures and tables. J.M. wrote the manuscript, with contributions from R.M. and M.A. participated in the design of the wet-lab experiment. J.M. and R.M. performed the analysis of the sequencing reads. M.A. and M.D. supervised the study. All authors read, revised, and approved the final manuscript.

## Competing interests

The authors declare no competing interests.

## Footnotes

a https://www.illumina.com/

b https://nanoporetech.com/

c https://www.agilent.com/

d https://www.dynegene.com/en/

e https://www.genscript.com/

## Notes

### Competing Interest Statement

The authors have declared no competing interest.

## References

1. Church, G. M., Gao, Y. & Kosuri, S. Next-generation digital information storage in DNA. Science 337, 1628–1628 (2012).

2. Grass, R. N., Heckel, R., Puddu, M., Paunescu, D. & Stark, W. J. Robust chemical preservation of digital information on dna in silica with error-correcting codes. Angewandte Chemie Int. Ed. 54, 2552–2555 (2015).

3. Organick, L. et al. Probing the physical limits of reliable dna data retrieval. Nat. communications 11, 616 (2020).

4. Zhirnov, V., Zadegan, R. M., Sandhu, G. S., Church, G. M. & Hughes, W. L. Nucleic acid memory. Nat. materials 15, 366–370 (2016).

5. Meiser, L. C. et al. Synthetic dna applications in information technology. Nat. communications 13, 352 (2022).

6. Coudy, D., Colotte, M., Luis, A., Tuffet, S. & Bonnet, J. Long term conservation of dna at ambient temperature. implications for dna data storage. PLoS One 16, e0259868 (2021).

7. Garibyan, L. & Avashia, N. Research techniques made simple: polymerase chain reaction (pcr). The J. investigative dermatology 133, e6 (2013).

8. Lehman, I. Discovery of dna polymerase. J. Biol. Chem. 278, 34733–34738 (2003).

9. Goldman, N. et al. Towards practical, high-capacity, low-maintenance information storage in synthesized dna. nature 494, 77–80 (2013).

10. Yazdi, S. H. T., Gabrys, R. & Milenkovic, O. Portable and error-free dna-based data storage. Sci. reports 7, 5011 (2017).

11. Pic, X. & Antonini, M. A constrained shannon-fano entropy coder for image storage in synthetic dna. In 2022 30th European Signal Processing Conference (EUSIPCO), 1367–1371 (IEEE, 2022).

12. Erlich, Y. & Zielinski, D. Dna fountain enables a robust and efficient storage architecture. science 355, 950–954 (2017).

13. Organick, L. et al. Random access in large-scale DNA data storage. Nat. biotechnology 36, 242–248 (2018).

14. Tomek, K. J. et al. Driving the scalability of dna-based information storage systems. ACS synthetic biology 8, 1241–1248 (2019).

15. Ross, M. G. et al. Characterizing and measuring bias in sequence data. Genome biology 14, R51 (2013).

16. Schwartz, J. J., Lee, C. & Shendure, J. Accurate gene synthesis with tag-directed retrieval of sequence-verified dna molecules. Nat. methods 9, 913–915 (2012).

17. Khabbaz, R., Mateos, J., Antonini, M. & Hanna, S. K. Dna-mgc+: A versatile codec for reliable and resource-efficient data storage on synthetic dna. arXiv preprint arXiv:2603.14527 (2026). Accepted for publication at Nature Communications Engineering 2026.

18. Cao, B. et al. Minimum free energy coding for dna storage. IEEE Transactions on Nanobioscience 20, 212–222 (2021).

19. Lazzarotto, D. et al. Overview of jpeg dna coding system for image storage on synthetic dna. In Applications of Digital Image Processing XLVIII, vol. 13605, 175–198 (SPIE, 2025).

20. Testolina, M. et al. JPEG AIC-3 Dataset: Towards Defining the High Quality to Nearly Visually Lossless Quality Range. In 2023 Fifteenth International Conference on Quality of Multimedia Experience (QoMEX), 55–60, DOI: 10.1109/QoMEX58391.2023.10178554 (2023).

21. JPEG Committee. JPEG DNA. https://jpeg.org/jpegdna/index.html.

22. ISO/IEC. ISO/IEC DIS 25508-1: Information technology — JPEG DNA media storage based on DNA — part 1: Core coding system. https://www.iso.org/standard/90579.html (2026). Draft International Standard.

23. Alakuijala, J., Sneyers, J., Versari, L. & Wassenberg, J. Jpeg white paper: Jpeg xl image coding system. Techn. Ber. ISO/IEC JTC 1, 1–3 (2023).

24. Luby, M., Shokrollahi, A., Watson, M. & Stockhammer, T. Raptor forward error correction scheme for object delivery. Tech. Rep. RFC 5053, Internet Engineering Task Force (2007).

25. Mateos, J., Lavenier, D., Dimopoulou, M., Genot, A. & Antonini, M. Primer design for DNA storage random access. In CORESA 2024 - 23ème conférence sur COmpression et REprésentation des Signaux Audiovisuels, 1–3 (Rennes, France, 2024).

26. Fornace, M. E., Porubsky, N. J. & Pierce, N. A. A unified dynamic programming framework for the analysis of interacting nucleic acid strands: Enhanced models, scalability, and speed. ACS Synth. Biol. 9, 2665–2678, DOI: 10.1021/acssynbio.9b00523 (2020).

27. Dirks, R. M., Bois, J. S., Schaeffer, J. M., Winfree, E. & Pierce, N. A. Thermodynamic analysis of interacting nucleic acid strands. SIAM Rev. 49, 65–88, DOI: 10.1137/060651100 (2007).

28. Li, H. Minimap2: pairwise alignment for nucleotide sequences. Bioinformatics 34, 3094–3100 (2018).

29. Ewing, B. & Green, P. Base-calling of automated sequencer traces using Phred. II. error probabilities. Genome Res. 8, 186–194, DOI: 10.1101/gr.8.3.186 (1998).

30. Martin, M. Cutadapt removes adapter sequences from high-throughput sequencing reads. EMBnet. journal 17, 10–12 (2011).

31. Fu, L., Niu, B., Zhu, Z., Wu, S. & Li, W. CD-HIT: accelerated for clustering the next-generation sequencing data. Bioinformatics 28, 3150–3152 (2012).

32. Mateos, J., Moreau, R., Dimopoulou, M. & Antonini, M. Jpeg dna thermodynamically constrained encoding subset. Zenodo, DOI: 10.5281/zenodo.22712282 (2026).

